# Genome-wide Association Study of Pancreatic Fat: The Multiethnic Cohort Adiposity Phenotype Study

**DOI:** 10.1101/2021.03.23.436581

**Authors:** Samantha A Streicher, Unhee Lim, S. Lani Park, Yuqing Li, Xin Sheng, Victor Hom, Lucy Xia, Loreall Pooler, John Shepherd, Lenora WM Loo, Burcu F Darst, Heather M Highland, Linda M Polfus, David Bogumil, Thomas Ernst, Steven Buchthal, Adrian A Franke, Veronica Wendy Setiawan, Maarit Tiirikainen, Lynne R Wilkens, Christopher A Haiman, Daniel O Stram, Iona Cheng, Loïc Le Marchand

## Abstract

Several studies have found associations between higher pancreatic fat content and adverse health outcomes, such as diabetes and the metabolic syndrome, but investigations into the genetic contributions to pancreatic fat are limited. This genome-wide association study, comprised of 804 participants with MRI-assessed pancreatic fat measurements, was conducted in the ethnically diverse Multiethnic Cohort-Adiposity Phenotype Study (MEC-APS). Two genetic variants reaching genome-wide significance, rs73449607 on chromosome 13q21.2 (Beta = −0.67, P = 4.50×10^-8^) and rs7996760 on chromosome 6q14 (Beta = −0.90, P = 4.91×10^-8^) were associated with percent pancreatic fat on the log scale. Rs73449607 was most common in the African American population (13%) and rs79967607 was most common in the European American population (6%). Rs73449607 was also suggestively associated with lower risk of type 2 diabetes (OR = 0.95, 95% CI = 0.89-1.00, P = 0.047) in the Population Architecture Genomics and Epidemiology (PAGE) Study and the DIAbetes Genetics Replication and Meta-analysis (DIAGRAM), which included substantial numbers of non-European ancestry participants (53,102 cases and 193,679 controls). Rs73449607 is located in an intergenic region between *GSX1* and *PLUT*, and rs79967607 is in intron 1 of *EPM2A*. *PLUT*, *a linkRNA*, regulates transcription of an adjacent gene, *PDX1*, that controls beta-cell function in the mature pancreas, and *EPM2A* encodes the protein laforin, which plays a critical role in regulating glycogen production. If validated, these variants may suggest a genetic component for pancreatic fat and a common etiologic link between pancreatic fat and type 2 diabetes.

## Introduction

Pancreatic fat accumulation (also referred to as pancreatic steatosis or pancreatic lipomatosis) was first described in the 1920s. Due to difficulties in obtaining pancreatic specimens, the effect of pancreatic fat on health outcomes began only to be explored over the last decade when new imaging modalities including ultrasonography (US), computed tomography (CT), and magnetic resonance imaging (MRI) have allowed researchers to non-invasively visualize internal organs [1–4]. Although diagnostic error rates from imaging machine variability and operator errors are factors for all types of data collection, MRI has emerged as the most sensitive non-invasive method for detection and quantification of pancreatic fat [3–5].

Pancreatic fat accumulation has been examined mainly in European populations [6]. In the few studies that have included non-Europeans, the amount of pancreatic fat accumulation was seen to vary by racial/ethnic groups [7, 8]. In a small study of overweight self-reported African American and Hispanic participants, African American participants were found to have a lower MRI-assessed mean percent pancreatic fat compared to Hispanic participants (P<0.0001) [7]. Additionally, in a small study of mildly obese self-reported African American, Hispanic, and white participants, magnetic resonance spectroscopy (MRS)-assessed mean pancreatic triglyceride levels were significantly lower in Black participants compared to Hispanic and white participants (P=0.006) [8].

Recently, Singh and colleagues (2017) conducted a meta-analysis in European American populations on the association between non-alcoholic fatty pancreas disease (NAFPD) and common metabolic diseases [6]. NAFPD (defined as >6.2% pancreatic fat in individuals consuming non-excessive amounts of alcohol) was found to be strongly associated with diabetes (risk ratio (RR)= 2.08, 95% confidence interval (95% CI): 1.44-3.00), the metabolic syndrome (RR=2.37, 95% CI=2.07-2.71), non-alcoholic fatty liver disease (NAFLD) (RR=2.67, 95% CI: 2.00-3.56), and hypertension (RR=1.67, 95% CI: 1.32-2.10) [6], after adjustment for possible confounding variables. While the pathophysiology of pancreatic fat remains to be fully elucidated, there is evidence suggesting that accumulation of pancreatic fat can occur from either the death of pancreatic acinar cells followed by adipocyte replacement, or by adipocyte infiltration of the pancreas caused by obesity [9].

Research has shown that the process of pancreatic fat infiltration and associated adverse health outcomes may be partially reversible through diet, exercise, and/or bariatric surgery [10–12], including a study that revealed reduction in pancreatic triglyceride levels only in type 2 diabetes (T2D) patients and not in normal glucose tolerance patients after bariatric surgery [12]. This finding, along with the research showing varying amounts of pancreatic fat by race/ethnicity, further raise the possibility of a genetic component for pancreatic fat accumulation that has yet to be explored. Therefore, in this study, we conducted a GWAS of pancreatic fat evaluated by MRI in the Multiethnic Cohort-Adiposity Phenotype Study (MEC-APS) and examined two identified genome-wide significant variants for association with obesity-related biomarkers in MEC-APS, and with T2D in independent populations.

## Results

The GWAS study population consisted of 804 MEC-APS study participants, including 144 African Americans, 129 European Americans, 206 Japanese Americans, 187 Latinos, and 138 Native Hawaiians (**Table 1**). Median overall age at clinic visit was 69.1 years (**Table 1**). Study participants in the lowest quartile (0.74-1.91%) of percent pancreatic fat were more likely to be African American, have the lowest mean BMI, total fat mass, visceral fat area, subcutaneous fat area, and percent liver fat, compared to participants in the three higher quartiles. Participants who were in the highest quartile (5.11-26.6%) of percent pancreatic fat were more likely to be Japanese American, have the highest mean BMI, total fat mass, visceral fat area, subcutaneous fat area, and percent liver fat compared to participants in the three lower quartiles (**Table 1**).

**Table 1.** Descriptive characteristics of MEC-APS subject by quartiles of percent pancreatic fat (N=804)^a^

|  | **Overall**<br>**(N=804)** | **Quartile 1**<br>**0.74-1.91 Percent**<br>**Pancreas Fat (n=201)** | **Quartile 2**<br>**1.92-3.22 Percent**<br>**Pancreas Fat (n=201)** | **Quartile 3**<br>**3.23-5.10 Percent**<br>**Pancreas Fat (n=201)** | **Quartile 4**<br>**5.11-26.6 Percent**<br>**Pancreas Fat (n=201)** |
| --- | --- | --- | --- | --- | --- |
| Age at clinic visit, years | 69.1 (67.1, 71.1) | 68.5 (67.1, 70.8) | 69.0 (67.1, 71.1) | 69.5 (67.0, 70.9) | 69.9 (67.4, 71.4) |
| Sex, n (%) |  |  |  |  |  |
| Men | 421 (52%) | 97 (49%) | 107 (53%) | 107 (53%) | 111 (55%) |
| Women | 383 (48%) | 103 (51%) | 94 (47%) | 94 (47%) | 90 (45%) |
| Race/ethnicity, n (%) |  |  |  |  |  |
| African American | 144 (17.9%) | 54 (27%) | 32 (16%) | 28 (14%) | 26 (13%) |
| European American | 129 (16%) | 28 (14%) | 24 (12%) | 40 (17%) | 40 (20%) |
| Japanese American | 206 (27%) | 46 (23%) | 62 (31%) | 36 (18%) | 60 (30%) |
| Latino | 187 (23.3%) | 46 (23%) | 52 (26%) | 60 (30%) | 28 (14%) |
| Native Hawaiian | 138 (17.2%) | 24 (12%) | 28 (14%) | 40 (20%) | 44 (22%) |
| Body mass index, kg/m<sup>2</sup> | 27.7 (24.9, 30.8) | 25.5 (23.4, 29.0) | 27.9 (24.8, 30.3) | 27.9 (25.6, 30.9) | 29.2 (26.9, 32.2) |
| Total fat mass, kg | 24.6 (19.5, 30.1) | 22.3 (17.7, 27.7) | 24.3 (19.5, 30.6) | 26.2 (20.3, 31.3) | 26.4 (21.9, 30.9) |
| Visceral fat area (L1-L5), cm<sup>2</sup> | 24.3 (19.4, 29.9) | 128.6 (88.0, 176.6) | 150.0 (118.8, 200.2) | 172.3 (132.7, 216.1) | 199.1 (147.9, 251.5) |
| Subcutaneous fat area (L1-L5), cm<sup>2</sup> | 33.0 (25.9, 40.6) | 179.3 (142.5, 242.7) | 211.0 (155.6, 285.9) | 222.6 (168.9, 298.1) | 239.3 (177.9, 300.0) |
| Liver fat, % | 4.3 (2.9, 7.5) | 3.6 (2.5, 5.6) | 4.4 (2.9, 8.4) | 5.0 (3.2, 8.7) | 5.2 (3.2, 8.2) |

<sup>a</sup>Count (percentage) of categorical variables and median (interquartile range) of continuous variables are presented across quartiles of percent pancreatic fat.

Overall, percent pancreatic fat had weak to moderate linear correlations with total fat mass (r=0.22), visceral fat area (r=0.34), subcutaneous fat area (r=0.20), and percent liver fat (r=0.17) (**Supplementary Table 2**). These correlations differed slightly by race/ethnicity, but remained weak to moderate.

In the MEC-APS, two loci were associated significantly with pancreatic fat at the genome-wide level: rs73449607 on chromosome 13q21.2 in an intergenic region between *GSX1* (GS Homeobox 1) and *PLUT* (*PDX1* associated long non-coding RNA, upregulator of transcription) and rs79967607 on 6q14 in intron 1 of the *EPM2A* gene (**Figures 1 and 2**, **Table 2**). The T allele of rs73449607 on chromosome 13q21.2 was associated with a 0.49-fold (95% CI = 0.40-0.65) decrease in geometric mean percent pancreatic fat (Beta = −0.67, P = 4.50×10^-8^), independent of age, sex, and principal components (**Table 2**). The geometric mean percent pancreatic fat for subjects who were homozygous recessive (TT), heterozygous (TC or CT), or homozygous dominant (CC) at rs73449607 was 0.58, 1.51, or 3.05, respectively. The T allele of rs73449607 was also associated with a non-significant decrease in the odds of NAFPD (OR = 0.15; 95% CI = 0.02-1.25) (**Supplementary Tables 3**). The rs73449607 association with pancreatic fat remained suggestive with additional adjustment for total fat mass (Beta = −0.27, P = 1.62×10^-7^) (**Supplementary Table 4**). While rs73449607 had a strong association with percent pancreatic fat, weaker associations existed with total fat mass (Beta = −0.09, P = 0.05), visceral fat area (Beta = −0.13, P = 0.04), subcutaneous fat area (Beta = −0.13, P = 0.02), and percent liver fat (Beta = −0.12, P = 0.26) (**Supplementary Table 5**). The association between rs73449607 and percent pancreatic fat appeared to have a larger effect and smaller P-value in men compared to women, but the interaction between rs73449607 and sex was not statistically significant (P=0.12) (**Table 2**). Overall, rs73449607 explained 5.3% of the variance in percent pancreatic fat. The T allele of rs73449607 was most frequent in African Americans (13%), present at low frequency in Latinos (1.1%), rare in Japanese Americans (0.2%), and not observed in Native Hawaiians or European Americans (**Table 3**). The most significant association across race/ethnicity between rs73449607 and percent pancreatic fat was in African Americans (Beta = −0.62; P = 9.60 × 10^-7^) with consistent effect estimates and directions of associations in the other non-monomorphic populations (Latinos and Japanese Americans) (**Table 3**). The interaction between the effect of rs73449607 and race/ethnicity did not reach statistical significance (P=0.28) (**Table 3**). In the African American population, rs73449607 explained 14.3% of the variance in percent pancreatic fat. Overall, in PAGE/DIAGRAM, rs73449607 also showed a nominally significant association with decreased risk of T2D (OR = 0.95; 95% CI = 0.89-1.00; P = 0.047) (**Table 4**). This association was driven by the African American (OR = 0.96; 95% CI = 0.90-1.02; P = 0.20) and Hispanic (OR = 0.86; 95% CI = 0.74-1.00; P = 0.047) populations (**Supplementary Table 6**). Of the 11 obesity-related circulating biomarkers examined in MEC-APS participants, the T allele of rs73449607 was associated with a 1.25-fold increase (Beta = 0.22; P = 1.2×10^-4^) in geometric mean for SHBG (**Table 5**). No association was found with other biomarkers, including glucose, insulin, or HOMA-IR (**Table 5**).

**Figure 1.**
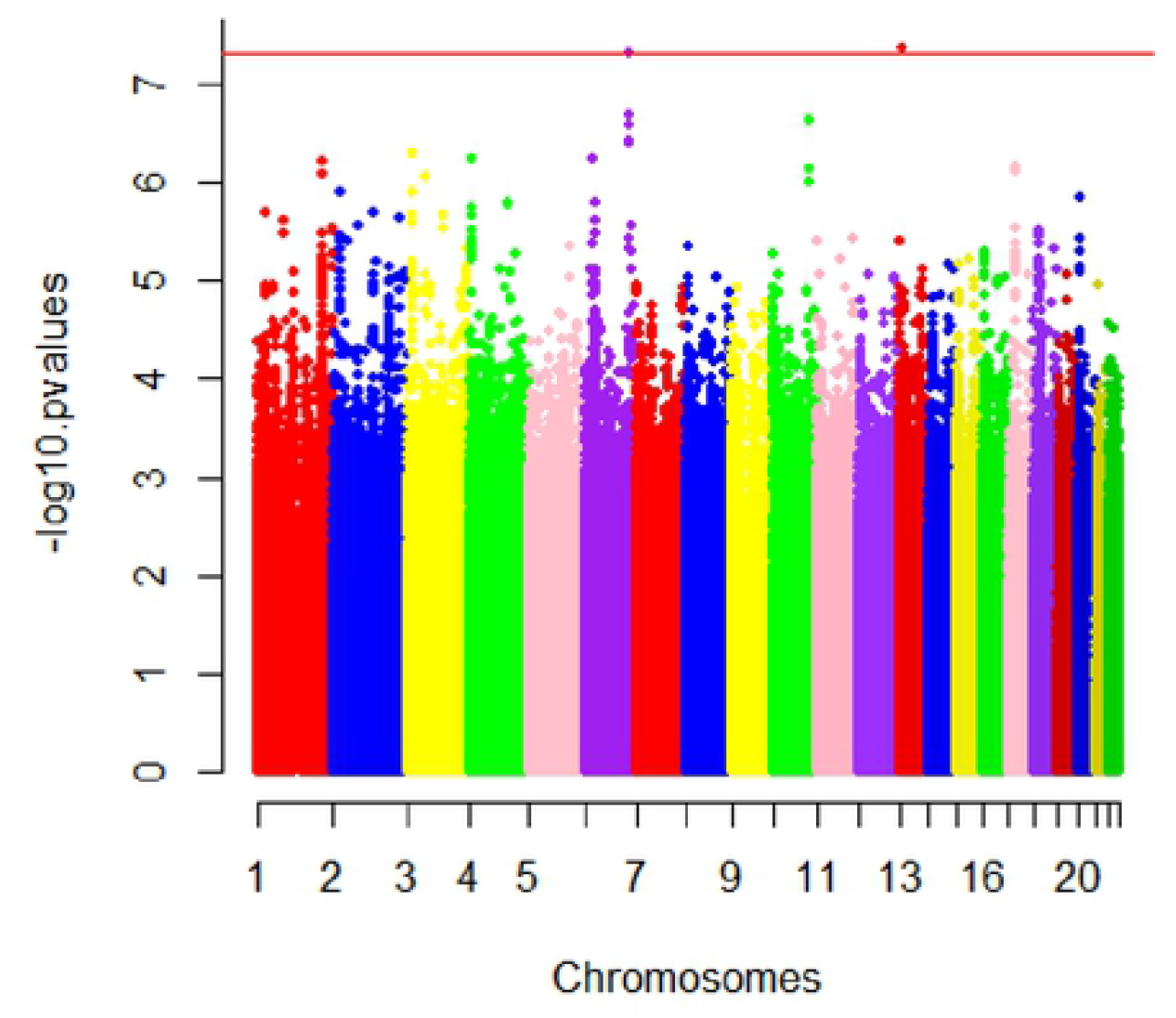
Manhattan plot of SNP P-values from the pancreas fat genome-wide association study in the Multiethnic Cohort-Adiposity Phenotype Study (MEC-APS). The Y-axis shows the negative base ten logarithm of the P-values and the X-axis shows the chromosomes. The genome-wide significance threshold, P<5×10^-8^, is shown in red.

**Figure 2.**
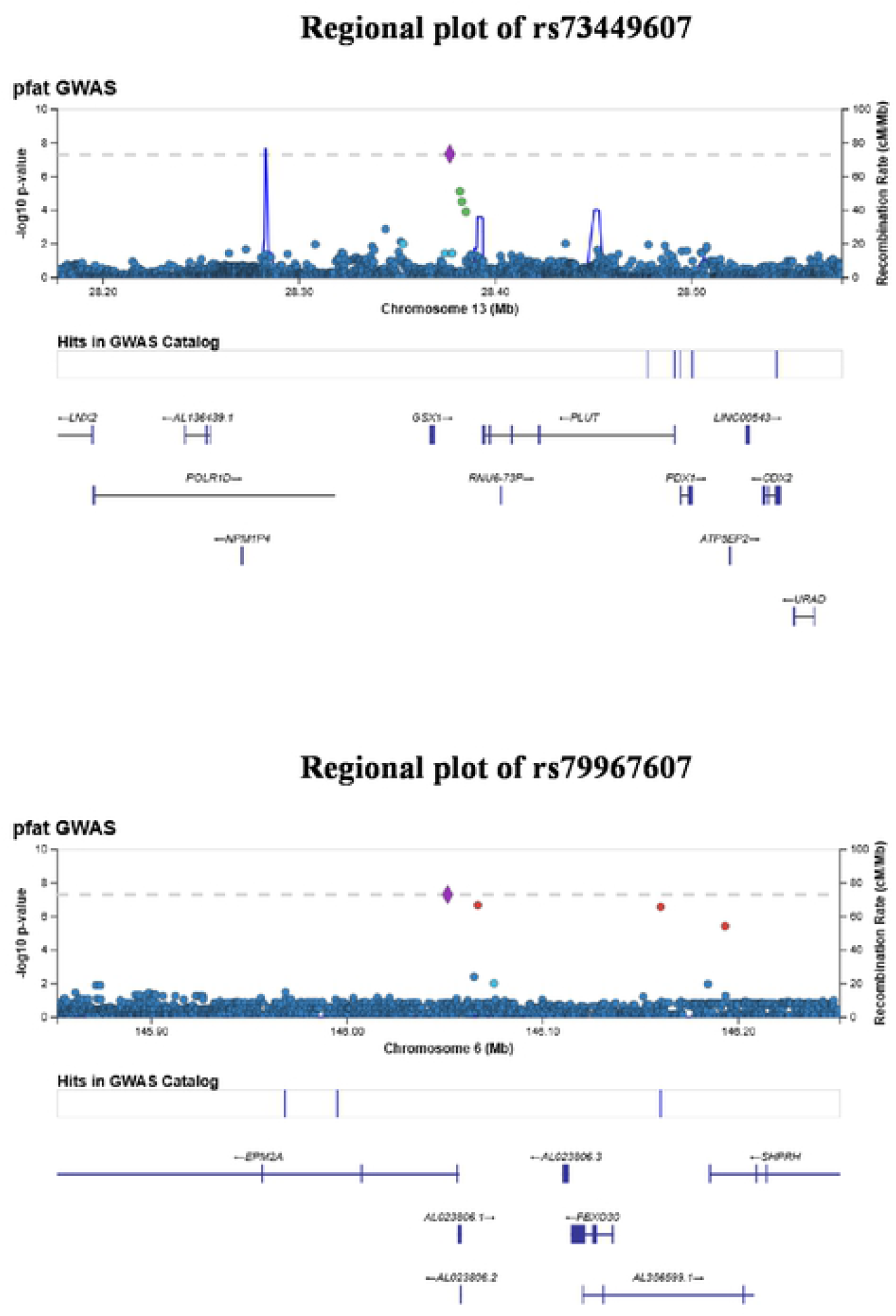
Regional plots of SNP P-values in a +/-200 kb window around rs73449607 and rs79967607. The X-axis shows the chromosome and physical location (Mb), the left Y-axis shows the negative base ten logarithm of the P-values, and the right Y-axis shows recombination activity (cM/Mb) as a blue line. Positions, recombination rates, and gene annotations are according to NCBI’s build 37 (hg 19) and the 1000 Genomes Project Phase 3 multiethnic data set.

**Table 2.** Two genetic variants associated with percent pancreatic fat in the MEC-APS (P<5 x 10^-8^) and median (interquartile range) of percent pancreatic fat, overall and by sex

|  |  |  |  |  |  | Overall (N=804) |  |  |  | Male (n=421) |  |  |  | Female (n=383) |  |  |  | P-het |
| --- | --- | --- | --- | --- | --- | --- | --- | --- | --- | --- | --- | --- | --- | --- | --- | --- | --- | --- |
| SNP | Chr<sup>d</sup> | Position<sup>e</sup> | Imputed Info score | Ref Allele | Effect Allele | EAf<sup>f</sup> | Beta<sup>g</sup> | SE | P-value | EAf<sup>c</sup> | Beta<sup>d</sup> | SE | P-value | EAf<sup>c</sup> | Beta<sup>d</sup> | SE | P-value |  |
| rs73449607<sup>a,b</sup> | 13 | 28376759 | 0.91 | C | T | 0.026 | -0.67 | 0.12 | 4.50x10<sup>-8</sup> | 0.021 | -1.00 | 0.19 | 5.65x10<sup>-7</sup> | 0.032 | -0.45 | 0.15 | 0.0034 | 0.12 |
| rs79967607<sup>a,c</sup> | 6 | 146051328 | 0.86 | T | G | 0.016 | -0.90 | 0.16 | 4.91x10<sup>-8</sup> | 0.015 | -0.89 | 0.24 | 2.64x10<sup>-4</sup> | 0.018 | -0.89 | 0.22 | 7.33x10<sup>-5</sup> | 0.55 |
| Percent pancreatic fat |  |  |  |  |  | 3.2 (1.9-5.1) |  |  |  | 3.3 (1.9-4.7) |  |  |  | 3.1 (1.8-4.8) |  |  |  |  |

<sup>a</sup>Adjusted for age, sex, and principal components 1-4. <sup>b</sup>For rs73449607, in the overall, male, and female population there were approximately 42, 18, and 25 T alleles, respectively. <sup>c</sup>For rs79967607, in the overall, male, and female population there were 26, 13, and 14 G alleles, respectively. <sup>d</sup>Chr, chromosome. <sup>e</sup>Position according to NCBI build37. <sup>f</sup>EAf, Effect allele frequency, <sup>g</sup>Log unit change per allele increase.

**Table 3.**
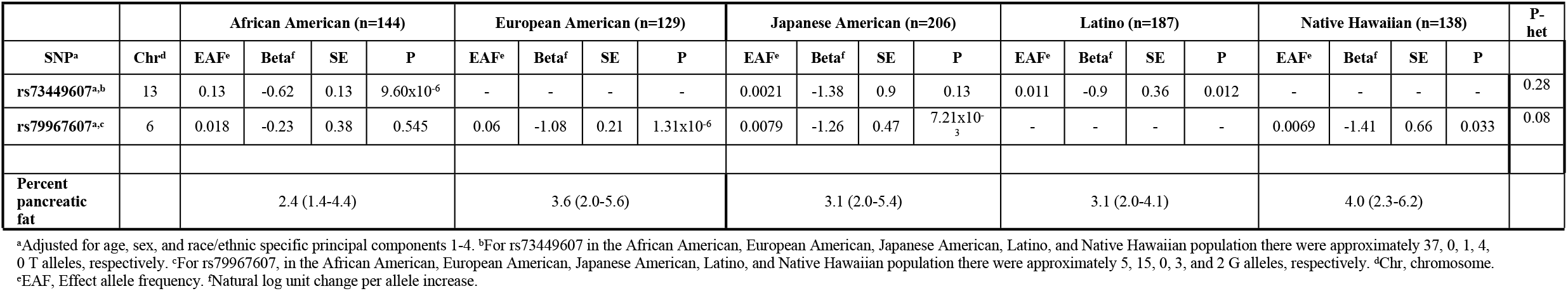
The association between rs73449607 or rs79967607 and pancreatic fat in the MEC-APS and median of percent pancreatic fat (interquartile range), by race/ethnicity

|  |  | African American (n=144) |  |  |  | European American (n=129) |  |  |  | Japanese American (n=206) |  |  |  | Latino (n=187) |  |  |  | Native Hawaiian (n=138) |  |  |  | P-het |
| --- | --- | --- | --- | --- | --- | --- | --- | --- | --- | --- | --- | --- | --- | --- | --- | --- | --- | --- | --- | --- | --- | --- |
| SNP<sup>a</sup> | Chr<sup>d</sup> | EAFe | Beta<sup>f</sup> | SE | P | EAFe | Beta<sup>f</sup> | SE | P | EAFe | Beta<sup>f</sup> | SE | P | EAFe | Beta<sup>f</sup> | SE | P | EAFe | Beta<sup>f</sup> | SE | P |  |
| rs73449607<sup>a,b</sup> | 13 | 0.13 | -0.62 | 0.13 | 9.60x10<sup>-6</sup> | - | - | - | - | 0.0021 | -1.38 | 0.9 | 0.13 | 0.011 | -0.9 | 0.36 | 0.012 | - | - | - | - | 0.28 |
| rs79967607<sup>a,c</sup> | 6 | 0.018 | -0.23 | 0.38 | 0.545 | 0.06 | -1.08 | 0.21 | 1.31x10<sup>-6</sup> | 0.0079 | -1.26 | 0.47 | 7.21x10<sup>-3</sup> | - | - | - | - | 0.0069 | -1.41 | 0.66 | 0.033 | 0.08 |
| Percent pancreatic fat |  | 2.4 (1.4-4.4) |  |  |  | 3.6 (2.0-5.6) |  |  |  | 3.1 (2.0-5.4) |  |  |  | 3.1 (2.0-4.1) |  |  |  | 4.0 (2.3-6.2) |  |  |  |  |

<sup>a</sup>Adjusted for age, sex, and race/ethnic specific principal components 1-4. <sup>b</sup>For rs73449607 in the African American, European American, Japanese American, Latino, and Native Hawaiian population there were approximately 37, 0, 1, 4, 0 T alleles, respectively. <sup>c</sup>For rs79967607, in the African American, European American, Japanese American, Latino, and Native Hawaiian population there were approximately 5, 15, 0, 3, and 2 G alleles, respectively. <sup>d</sup>Chr, chromosome. <sup>e</sup>EAFe, Effect allele frequency. <sup>f</sup>Natural log unit change per allele increase.

**Table 4.** The association between rs73449607 or rs79967607 and Type 2 Diabetes in the Population Architecture Genomics and Epidemiology/DIAbetes Genetics Replication and Meta-analysis (PAGE/DIAGRAM) study

| SNP | Chr<sup>b</sup> | Position<sup>c</sup> | Imputed Info score<sup>d</sup> | Ref Allele | Effect Allele | Case EAF<sup>e,f</sup> | Control EAF<sup>e,f</sup> | OR (95% CI)<sup>g</sup> | P-value |
| --- | --- | --- | --- | --- | --- | --- | --- | --- | --- |
| rs73449607<sup>a</sup> | 13 | 28376759 | 0.72-0.98 | C | T | 0.0257 | 0.0143 | 0.95 (0.89, 1.00) | 0.0517 |
| rs79967607<sup>a</sup> | 6 | 146051328 | 0.81-0.97 | T | G | 0.0440 | 0.0487 | 0.96 (0.91, 1.01) | 0.146 |

<sup>a</sup>Adjusted for age, sex, BMI and principal components. <sup>b</sup>Chr, chromosome. <sup>c</sup>Position according to NCBI build37. <sup>d</sup>Imputation score is presented over a range from 24 different genotyping platforms. <sup>e</sup>EAF, Effect allele frequency. <sup>f</sup>Effect allele frequencies were calculated based on the PAGE MEGA array data for African, Hispanic, Asian, and Native Hawaiian populations and the WHIMS data for European populations. <sup>g</sup>OR, odds ratio; 95% CI, 95% Confidence Interval.

**Table 5.** The association between rs3449607 or rs79967697 and obesity-related biomarkers in the MEC-APS

| Variant | Biomarker | N | Beta<sup>b</sup> | SE | P-value |
| --- | --- | --- | --- | --- | --- |
| rs73449607<sup>a</sup> | HDL (mg/dL) | 1822 | 0.015 | 0.050 | 0.78 |
|  | LDL (mg/dL) | 1817 | -0.022 | 0.047 | 0.63 |
|  | Total Cholesterol (mg/dL) | 1823 | -0.0014 | 0.032 | 0.96 |
|  | Glucose (mg/dL) | 1821 | -0.011 | 0.027 | 0.68 |
|  | HOMA-beta (%) | 1810 | -0.12 | 0.099 | 0.24 |
|  | HOMA-IR | 1821 | -0.057 | 0.081 | 0.48 |
|  | CRP (mg/L) | 1823 | -0.051 | 0.020 | 0.80 |
|  | Insulin (microU/mL) | 1823 | -0.062 | 0.073 | 0.40 |
|  | SHBG (nmol/L) | 1816 | 0.22 | 0.057 | 1.25 x 10<sup>-4</sup> |
|  | Triglycerides (mg/dL) | 1823 | 0.053 | 0.053 | 0.57 |
|  | ALT (U/L) | 1823 | -0.054 | 0.056 | 0.34 |
| rs79967607<sup>a</sup> |  |  |  |  |  |
|  | HDL (mg/dL) | 1822 | 0.093 | 0.054 | 0.082 |
|  | LDL (mg/dL) | 1817 | 0.017 | 0.05 | 0.72 |
|  | Total Cholesterol (mg/dL) | 1823 | 0.033 | 0.17 | 0.33 |
|  | Glucose (mg/dL) | 1821 | 0.040 | 0.29 | 0.17 |
|  | HOMA-beta (%) | 1810 | -0.14 | 0.10 | 0.16 |
|  | HOMA-IR | 1821 | 0.019 | 0.086 | 0.82 |
|  | CRP (mg/L) | 1823 | 0.078 | 0.22 | 0.71 |
|  | Insulin (microU/mL) | 1823 | -0.021 | 0.078 | 0.79 |
|  | SHBG (nmol/L) | 1861 | 0.012 | 0.061 | 0.83 |
|  | Triglycerides (mg/dL) | 1823 | -0.031 | 0.57 | 0.59 |
|  | ALT (U/L) | 1823 | -0.0011 | 0.061 | 0.98 |

<sup>a</sup>Adjusted for age, sex, principal components 1-4, and total fat mass (kg). <sup>b</sup>Log unit change per allele increase.

The G allele of rs79967607 on chromosome 6q14 was associated with a 0.41-fold (95% CI = 0.29-0.56) decrease in geometric mean percent pancreatic fat (Beta = −0.90, P = 4.91×10^-8^) (**Table 2**). The geometric mean percent pancreatic fat for subjects who were heterozygous (GT or TG) or homozygous dominant (TT) at rs79967607 was 1.40 or 3.02, respectively. There was no participant homozygous recessive (GG) for rs79967607. The G allele of rs79967607 was also associated with a non-significant decrease in the odds of NAFPD (OR = 0.39; 95% CI = 0.09-1.69) (**Supplementary Tables 3**). The rs79967607 association with pancreatic fat remained suggestive after additional adjustment for total fat mass (Beta = −0.29, P = 2.81×10^-5^) (**Supplementary Tables 4**). While rs79967607 was strongly associated with percent pancreatic fat, the variant showed weaker associations with total fat mass (Beta = −0.12, P = 0.05), visceral fat area (Beta = −0.13, P = 0.04), subcutaneous fat area (Beta = −0.19, P = 0.01), and percent liver fat (Beta = −0.064, P = 0.64) (**Supplementary Table 5**). Overall, rs79967607 explained 3.6% of the variance in percent pancreatic fat. The G allele of rs79967607 was most frequent in European Americans (6%) followed by African Americans (2%), rare in Latinos (0.8%) and Native Hawaiians (0.8%), and not observed in Japanese Americans. For the association between rs79967607 and percent pancreatic fat, the most significant association was in European Americans (Beta = −1.08; P = 1.31 × 10^-6^) with consistent effect estimates and direction of associations in the other non-monomorphic populations (African Americans, Latinos, and Native Hawaiians) (**Table 3**). The test for interaction between rs79967607 and race/ethnicity was borderline significant (P=0.08) (**Table 3**). In the European American population, rs79967607 explained 11.9% of the variance in percent pancreatic fat. Overall, in PAGE/DIAGRAM, rs79967607 was not significantly associated with T2D (OR = 0.96; 95% CI = 0.91-1.01; P = 0.14) (**Table 4 and Supplementary Table 6**). Of the 11 obesity-related circulating biomarkers examined in MEC-APS participants, the G allele of rs79967607 was suggestively associated with a 1.10-fold increase (Beta = 0.09; P = 0.08) in the geometric mean of HDL (**Table 5**).

## Discussion

In our GWAS of pancreatic fat in a racially/ethnically diverse population, we observed genome-wide significant associations with percent pancreatic fat with rs73449607, a variant in an intergenic region on chromosome 13q21.2 and with rs79967607, a variant in intron 1 of *EPM2A* on chromosome 6q14. Both variants appear to be specific for pancreatic fat since they were only weakly associated with other total and ectopic adiposity phenotypes, all quantified using state-of-the-art imaging methods. Imputation quality for rs73449607 and rs79967607 was high, and estimates and P-values obtained from regressing percent pancreatic fat on retained imputed dosages (rs73449607: Beta = −0.67, P = 4.50×10^-8^ and rs79967607: Beta = −0.90, P = 4.91×10^-8^) were almost identical to the estimates and P-values obtained from regressing percent pancreatic fat on genotypes (rs73449607: Beta = −0.63, P = 1.74×10^-8^ and rs79967607: Beta = −0.84, P = 4.52×10^-8^), adjusted for age, sex, and principal components 1-4. Neither variant showed extreme Hardy-Weinberg departures.

The T allele of rs73449607 was associated with a 49% decrease in geometric mean percent pancreatic fat. While rs73449607 is closest to *GSX1* (~7.5kb downstream), it is also upstream of *PLUT* (also known as *HI-LNC71*, *PDX1-AS1*, and *PLUTO*) (~16kb upstream) and *PDX1* (pancreatic and duodenal homeobox) (~100kb upstream). There have been at least four genome-wide significant variants located in *PDX1* or *PLUT* found to be associated with fasting blood glucose or pancreatic cancer [13–16]. *PDX1* is an established regulator of early pancreatic development, has a role in differentiation of the exocrine pancreas, and regulates beta-cell function in the mature pancreas [17, 18]. Germline mutations in *PDX1* have been associated with agenesis of the pancreas and maturity onset diabetes of the young [19]. While less is known about *PLUT*, this gene has been shown to affect local 3D chromatin structure and transcription of *PDX1*, and both *PLUT* and *PDX1* have been shown to be downregulated in pancreatic islet cells obtained from individuals with T2D or impaired glucose tolerance [20]. In the overall analysis, rs73449607 was associated with decreased pancreatic fat content and was suggestively associated with a decreased risk of T2D. Notably, in African American participants, where the variant was most common, the T allele of rs73449607 was associated with a decreased amount of pancreatic fat and was nominally associated with a decreased risk of T2D. The variant rs73449607 was also associated with increased levels of the hormonal biomarker, SHBG. Interestingly, higher SHBG levels have been associated with a lower BMI and a decreased risk of T2D, but higher SHBG levels have also been found in advanced pancreatic cancer cases [21–23]. The association between rs73449607 and SHBG seemed to be independent of obesity since the effect estimate and P-value remained similar with (Beta = 0.09; P = 3.8×10^-4^) and without (Beta = 0.11; P = 3.3×10^-5^) adjustment for total fat mass.

The variant rs79967607 is located in intron 1 of *EPM2A*. The G allele of rs7996707 was associated with a 41% decrease in geometric mean percent pancreatic fat, after adjustment for age, sex, and principal components. The *EPM2A* gene encodes the protein laforin, which plays a critical role in regulating the production of glycogen [24]. When blood glucose rises, the pancreas responds by releasing insulin, which in turn lowers blood glucose levels by promoting the liver and muscles to take up glucose from the blood and store it as glycogen [24]. During times of physical exertion or fasting, glycogen can also be broken back down to glucose [24]. There is also some evidence that laforin may also act as a tumor suppressor protein [25]. Additionally, according to haploreg v4.1, rs79967607 is associated with enrichment of H3K4me1 epigenetic motifs in pancreatic islet cells [26]. In our analysis, rs79967607 was associated with pancreatic fat and not with T2D.

Although partial reversal of high pancreatic fat appears possible with weight loss, this study supports a genetic component to pancreatic fat deposition, which in turn, may influence other health outcomes [10–12, 27]. Our findings underscore the importance of conducting genetic analyses in multiethnic populations, as the significant variants varied in frequency across racial/ethnic groups, and rs73449607 was not associated with pancreatic fat in individuals of European ancestry [28]. Another strength of our analysis is that we used highly sensitive imaging methods to assess pancreatic and other ectopic fat amounts (MRI) and total fat mass (DXA), which provided the ability to test whether associations with pancreatic fat were independent of total fat mass.

In a preprint manuscript on bioRxiv, Liu and colleagues (2020) conducted GWASes of 11 MRI-assessed abdominal organ and adiposity measurements, including pancreatic volume and percent fat based on 30,000 UK Biobank participants of White British ancestry [29]. Regarding the two genome-wide significant variants in our study, rs73449607 was not observed in the European American population in MEC-APS and rs79967607 was not found to be genome-wide significantly associated with pancreatic fat in the UK Biobank population. However, 10 other significant variants were identified as genome-wide significant (P<5×10^-8^) in UK Biobank. Of these 10 significant variants [29], one variant showed a suggestive association with percent pancreatic fat in our MEC-APS study population (rs118005033: Beta = 0.10, P = 0.01), three variants were not associated with percent pancreatic fat (rs4733612: Beta = −0.07, P = 0.16; rs2270911: Beta = 0.04, P = 0.25; and rs13040225, Beta = 0.04, P = 0.27), and the remaining six variants were not in our final data set.

Although this is the first GWAS of pancreatic fat to be conducted in a multiethnic population, limitations to our study should be considered. First, due to the post-hoc measurements of pancreatic fat, only about half of the MRI scans had interpretable pancreas images. However, participant differences in interpretable and non-interpretable pancreas images were unlikely to explain our findings since sex and genetic ancestry (as principal components) were adjusted for in regression models and the genome-wide significant variants showed similar effect allele frequencies and similar or slightly stronger parameter estimates for participants with pancreatic fat data compared to all participants when other adiposity phenotypes were examined (**Supplementary Table 5**). Second, the total study population with MRI-assessed percent pancreatic fat was modest in size (N=804), and the study had limited statistical power to detect weak effects. Third, to our knowledge, the pancreas measurements on 30,000 participants of White British Ancestry from the UK Biobank is the only other comprehensive data set of participants with image-assessed pancreatic fat or biopsy and these are not accessible in the publically available data set, which makes replicating the association between our genome-wide significant variants and pancreatic fat challenging.

In summary, two variants, rs73449607 and rs79967607, were associated with percent pancreatic fat at the genome-wide significance level in our multiethnic GWAS. The variant rs73449607 also showed an association with blood levels of SHBG and a nominal association with T2D, while rs79967607 had a suggestive association with blood levels of HDL. Future large scale studies are needed to replicate these associations in large and diverse study populations and to identify additional variants associated with pancreatic fat. These variants, if validated, may point to biologic pathways for pancreatic fat and related health outcomes, such as T2D.

## Materials and Methods

### The Multiethnic Cohort-Adiposity Phenotype Study (MEC-APS)

The MEC was established in 1993-1996 to examine the association of lifestyle and genetics with cancer risk [30]. This prospective study has been following over 215,000 adult men and women living in Hawaii and California, predominately Los Angeles County. Participants are mostly from five main ethnic/racial groups (African American, Japanese American, Latino, Native Hawaiian, and European American) [30]. In 2013-2016, the MEC-APS was conducted to identify predictors of body fat distribution and obesity-related cancers, as described previously [31]. Briefly, this cross-sectional study recruited 1,861 healthy, not currently smoking, male and postmenopausal female MEC participants between 60-77 years of age, with no history of chronic hepatitis, and a body mass index (BMI) between 17.1-46.2 kg/m^2^. MEC participants were selected for the study using a stratified sampling by sex, race/ethnicity, and six BMI categories. All MEC-APS participants provided written informed consent and the study was approved by the institutional review boards (IRBs) at the University of Hawaii (CHS-#17200), University of Southern California (#HS-12-00623), and University of California, San Francisco (#17-23399) in agreement with the 1975 Helsinki Declaration. Study participants underwent an abdominal MRI and body composition assessment by whole-body dual energy X-ray absorptiometry (DXA), and completed blood collection, and self-administered questionnaires including a quantitative food-frequency questionnaire [31]. Seven participants were excluded after failing genotype quality control (QC) and 1,050 were excluded for missing percent pancreatic fat measurement. Since measurements of fat deposits in the pancreas were not originally included in the MEC-APS protocol, percent pancreatic fat measurements were ascertained post-hoc. Therefore, only about half of the MRI scans yielded interpretable pancreas images, due to differences in anatomical presentation (see below). Participants with interpretable pancreatic fat MRI images were more often men (P=0.04), Japanese Americans, Latinos, or Native Hawaiians (P<0.0001) and had greater visceral fat area (P=0.002) and percent liver fat (P<0.0001) compared to those with non-usable MRI (**SupplementaryTable 1**). There were no differences between the groups with and without valid pancreatic fat analysis by age (P=0.42), total adiposity (P=0.32), or subcutaneous fat area (P=0.09) (**Supplementary Table 1**). The final study population comprised 804 MEC-APS participants.

### Adiposity Measurements

The 3T MRI scans from a Siemens TIM Trio at UH and General Electric HDx at USC were used to quantify pancreatic fat, abdominal visceral and subcutaneous fat, and liver fat. Percent pancreatic fat was determined post-hoc from a series of axial triple gradient-echo Dixon-type MRI scans (10mm slices, no gap, TE=2.4, 3.7, and 5.0 ms, TR=160 ms, 25° flip angle) by analyzing in-phase, out-of-phase and in-phase signals in one or two circular regions of interest (ROI 15-20 cm^2^) in the pancreas, using all slices of images where a ROI could be captured while avoiding the folding of the pancreas. The Dixon protocol was applied to measure the proton density fat fraction (PDFF) of the liver and pancreas since it has shown high accuracy when compared to histologic fat fraction. It has also shown a high correlation with MR spectroscopy but has a shorter acquisition and processing time and a significantly higher sensitivity over ultrasound or computed tomography methods [32, 33]. Additional details regarding the protocol, as well as measurement of visceral fat area, subcutaneous fat area, and percent liver fat were previously published by Lim and colleagues (2019) [31]. NAFPD (188 cases and 549 controls) was defined as pancreatic fat >5% for participants with no excessive alcohol consumption (defined as >30 g/day of alcohol in men and >20 g/day of alcohol in women) in the past year [31, 34]. Total fat mass (kg) was measured by whole-body DXA scan (Hologic Discovery A densitometer, Bedford, MA) [35].

### Obesity -related biomarkers

Selected blood biomarkers were chosen for their reported associations with obesity-caused metabolic, hormonal, and inflammation dysfunctions [36]. Fasting blood samples were collected at the time of body composition measurement, processed into components, and stored at −80°C [36]. Concentrations of biomarkers (high density lipoprotein (HDL) (mg/dL) (N=1822), low density lipoprotein (LDL) (mg/dL) (N=1817), total cholesterol (mg/dL) (N=1823), glucose (mg/dL) (N=1821), homeostasis model assessment (HOMA)-beta (%) (N=1810), HOMA-insulin resistance (IR) (%) (N=1821), C-reactive protein (CRP) (mg/dL) (N=1823), insulin (microU/mL) (N=1823), sex hormone binding globulin (SHBG) (nmol/L) (N=1816), triglycerides (mg/dL) (N=1823), and alanine aminotransferase (ALT) (U/L) (N=1823) were measured in blood samples from plasma or serum: detailed assay protocols and good reproducibility have been reported previously [36]. HOMA-IR and HOMA-beta were derived from fasting glucose and insulin values [36–38]. LDL cholesterol was derived from the Friedewald equation using total cholesterol and HDL cholesterol concentrations and a valid range of triglyceride concentrations [39].

### Genotyping, Quality Control, and Imputation

Genotyping and imputation for the MEC-APS participants have been described previously [34]. Briefly, DNA extraction from buffy coat was performed using the Qiagen QIAMP DNA kit (Qiagen Inc., Valencia, CA). DNA samples were genotyped on the Illumina expanded multi-ethnic genotyping array (MEGA^EX^) platform, which to date provides the largest coverage of variants across the genome for diverse ancestral populations [40]. Variants with a call rate <95%, variants with a replicate concordance <100% based on 39 QC replicate samples, or variants with poor clustering after visual inspection were removed. Prior to imputation, monomorphic variants, variants with call rate <98%, variants with estimated minor allele frequency that deviated by ≥20% in comparison to the corresponding ancestral group in the 1000 Genomes Project Phase 3, discordance in reported vs. genotyped sex, and insertions/deletions which are not included in the Haplotype Reference Consortium (HRC), were removed. From an initial 2,036,060 genotyped variants, 1,417,570 were available for imputation. Phasing using Eagle v2.4 and genotype imputation using Minimac v4 were performed on the University of Michigan Imputation Server with the HRC vr1.1 2016 reference panel [41, 42]. After genotype imputation for MEC-APS participants, variants with an imputation quality score of < 0.4, multiallelic variants, variants with MAF<0.01, or monomorphic variants in either NAFPD cases or controls, were excluded from all subsequent analyses. In total, 9,542,479 genotyped and imputed SNPs remained after post-imputation filtering. Principal components for ancestry adjustment were calculated with 91,762 post-QC genotyped linkage disequilibrium (LD) pruned SNPs using EIGENSOFT v7 [43].

### Population Architecture Genomics and Epidemiology (PAGE) Study/DIAbetes Genetics Replication and Meta-analysis (DIAGRAM)

The PAGE/DIAGRAM T2D GWAS meta-analysis has been described previously [44] and was used in this study to examine the association of our pancreatic fat GWAS hits and T2D. In brief, a total of 246,781 participants from 6 case-control studies included in PAGE (ARIC, BioME, CARDIA, MEC, SOL, and WHI) and 15 case-control studies included in DIAGRAM (deCODE, DGDG, DGI, EGCUT-370, EGCUT-OMNI, EPIC-InterAct, FHS, FUSION, GoDARTS, HPFS, KORAgen, NHS, PIVUS, RS-I, ULSAM, and WTCCC) were included in a GWAS meta-analysis. There were 8,591 T2D cases and 16,887 controls of African ancestry, 3,124 T2D cases and 4,313 controls of Asian ancestry, 9,913 T2D cases and 22,958 controls from Hispanic populations, 1,642 T2D cases and 2,152 controls of Native Hawaiian ancestry, and 29,832 T2D cases and 147,369 controls of European ancestry [44]. Twenty-seven MEC-APS T2D cases and 151 controls were also included in the PAGE/DIAGRAM study.

Descriptive characteristics were examined in the overall study population and by quartile of percent pancreatic fat (0.074-1.91%, 1.92-3.22%, 3.23-5.10%, and 5.11-26.6%). The chi-square test was used to compare categorical variables and the one-way analysis of variance (ANOVA) test was used to compare continuous variables using R v3.6.1.

Pearson’s correlations between log-transformed percent pancreatic fat and log-transformed total fat mass (N=793), visceral fat area (N=799), subcutaneous fat area (N=799), and percent liver fat (N=801) were calculated overall, and by race/ethnicity in R v3.6.1.

Variant (as imputed dosages) associations with percent pancreatic fat were estimated using linear regressions of log-transformed percent pancreatic fat, adjusted for age, sex, and main principal components 1-4 using additive genetic models, and then rerun with additional adjustment for total fat mass. SNP associations were considered statistically significant at the genome-wide significance threshold P<5×10^-8^. A quantile–quantile plot of GWAS P-values indicated appropriate control of type I errors, with a genomic inflation (λ) value of 1.03 (**Supplementary Figure 1**). Imputed dosages were converted to genotypes based on a hard call threshold of 0.49999, and geometric means of percent pancreatic fat was calculated for homozygous recessive, heterozygous, and homozygous dominant genotypes. Interactions between variants significantly associated with percent pancreatic fat and sex or race were also evaluated by adding interaction terms between the variant and sex or race/ethnicity to each model. Models were further stratified by sex (male, female) and self-reported race/ethnicity (African American, European American, Japanese American, Latino, Native Hawaiian), and adjusted for age, sex, and race or sex-specific principal components. All analyses were done in PLINK v2.0

Variants significantly associated with percent pancreatic fat were further assessed for association with total fat mass, visceral fat area, subcutaneous fat area, and percent liver fat in MEC-APS in order to examine whether they had a broader role in adiposity accumulation. Each log-transformed adiposity phenotype was regressed on the significant variant, adjusting for age, sex, and principal components 1-4 overall (N=1,825 for total fat mass, 1,787 for visceral fat area and subcutaneous fat area, and 1,775 for percent liver fat) and limited to participants with pancreatic fat data (N=793 for total fat mass, 799 for visceral fat area and subcutaneous fat area, and 1,775 for percent liver fat) using R v.3.6.1.

Variation in percent pancreatic fat (R^2^) explained by each genome-wide significant variant was calculated by 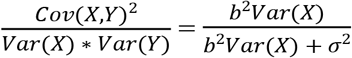 where *X* = the imputed dosage variable, *σ*^2^ = the variance of the residuals, and for a variant with the effect allele frequency *p*, *Var*(*X*) = 2*p*(1 – *p*), under the Hardy-Weinberg equilibrium (HWE) assumption.

Variants significantly associated with percent pancreatic fat were also assessed for relationships with NAFPD in MEC-APS (188 cases and 549 controls), with obesity-related biomarkers (HDL, LDL, total cholesterol, glucose, insulin, HOMA-beta, HOMA-IR, CRP, SHBG, triglycerides, and ALT) among over 1,800 MEC-APS participants (see above in *Obesity-related biomarkers* for exact number of participants analyzed for each biomarker), and with T2D among 53,102 cases and 193,679 controls in PAGE/DIAGRAM. Associations with NAFPD was assessed using logistic regression models adjusted for age, sex, total fat mass, and principal components 1-4. Associations with obesity-related biomarkers were assessed using linear regression models of log-transformed analytes adjusted for age, sex, total fat mass, and principal components 1-4. Both NAFPD and obesity-related biomarkers regression models were run in PLINK v2.0. Associations with T2D were assessed with unconditional logistic regression models adjusted for age, sex, body mass index, and principal components. Every racial/ethnic population within each T2D study was analyzed separately. Racial/ethnic population-specific estimates were obtained by combining per-allele odds ratios and standard errors across studies for each racial/ethnic population. Overall estimates were obtained by combining per-allele odds ratios and standard errors first across racial/ethnic populations within each study and then by combining per-allele odds ratios and standard errors across each study. Both racial/ethnic population-specific estimates and overall estimates were obtained using fixed-effects inverse-variance weighted meta-analyses, as implemented in METAL [44, 45].

## Acknowledgements

The PAGE consortium thanks the staff and participants of all PAGE studies for their important contributions. The complete list of PAGE members can be found at http://www.pagestudy.org.

## References

1. Shaefer J. The normal weight of the pancreas in the adult human being: A biometric study. The Anatomical Record. 1926;32:119–32.

2. Ogilvie R. The islands of langerhans in 19 cases of obesity. The Journal of Pathology and Bacteriology. 1933;37:473–81.

3. Dite P, Blaho M, Bojkova M, Jabandziev P, Kunovsky L. Nonalcoholic fatty pancreas disease: Clinical consequences. Dig Dis. 2020;38:143–9.

4. Yu TY, Wang CY. Impact of non-alcoholic fatty pancreas disease on glucose metabolism. J Diabetes Investig. 2017;8:735–47.

5. Kato S, Iwasaki A, Kurita Y, Arimoto J, Yamamoto T, Hasegawa S, et al. Three-dimensional analysis of pancreatic fat by fat-water magnetic resonance imaging provides detailed characterization of pancreatic steatosis with improved reproducibility. PLoS One. 2019;14:e0224921.

6. Singh RG, Yoon HD, Wu LM, Lu J, Plank LD, Petrov MS. Ectopic fat accumulation in the pancreas and its clinical relevance: A systematic review, meta-analysis, and meta-regression. Metabolism. 2017;69:1–13.

7. Le KA, Ventura EE, Fisher JQ, Davis JN, Weigensberg MJ, Punyanitya M, et al. Ethnic differences in pancreatic fat accumulation and its relationship with other fat depots and inflammatory markers. Diabetes Care. 2011;34:485–90.

8. Szczepaniak LS, Victor RG, Mathur R, Nelson MD, Szczepaniak EW, Tyer N, et al. Pancreatic steatosis and its relationship to beta-cell dysfunction in humans: racial and ethnic variations. Diabetes Care. 2012;35:2377–83.

9. MM S, EJM vG. The clinical sigificance of pancreatic steatosis. Nat Rev Gastroenterol Hepatol. 2011;8:169–77.

10. Lim EL, Hollingsworth KG, Aribisala BS, Chen MJ, Mathers JC, Taylor R. Reversal of type 2 diabetes: normalisation of beta cell function in association with decreased pancreas and liver triacylglycerol. Diabetologia. 2011;54:2506–14.

11. Taylor R. Type 2 diabetes: etiology and reversibility. Diabetes Care. 2013;36:1047–55.

12. Steven S, Hollingsworth KG, Small PK, Woodcock SA, Pucci A, Aribisala B, et al. Weight loss decreases excess pancreatic triacylglycerol specifically in type 2 diabetes. Diabetes Care. 2016;39:158–65.

13. Manning AK, Hivert MF, Scott RA, Grimsby JL, Bouatia-Naji N, Chen H, et al. A genome-wide approach accounting for body mass index identifies genetic variants influencing fasting glycemic traits and insulin resistance. Nat Genet. 2012;44:659–69.

14. Wolpin BM, Rizzato C, Kraft P, Kooperberg C, Petersen GM, Wang Z, et al. Genome-wide association study identifies multiple susceptibility loci for pancreatic cancer. Nat Genet. 2014;46:994–1000.

15. Nagy R, Boutin TS, Marten J, Huffman JE, Kerr SM, Campbell A, et al. Exploration of haplotype research consortium imputation for genome-wide association studies in 20,032 Generation Scotland participants. Genome Med. 2017;9:23–36.

16. Kanai M, Akiyama M, Takahashi A, Matoba N, Momozawa Y, Ikeda M, et al. Genetic analysis of quantitative traits in the Japanese population links cell types to complex human diseases. Nat Genet. 2018;50:390–400.

17. Stoffers D, Zinkin N, Stanojevic V, Clarke W, Habener J. Pancreatic agenesis attributable to a single nucleotide deletion in the human IPF1 gene coding sequence. Nature Genetics. 1997;15:106–10.

18. MacDonald RJ, Swift GH, Real FX. Transcriptional control of acinar development and homeostasis. Prog Mol Biol Transl Sci. 2010;97:1–40.

19. Vaxillaire M, Bonnefond A, Froguel P. The lessons of early-onset monogenic diabetes for the understanding of diabetes pathogenesis. Best Pract Res Clin Endocrinol Metab. 2012;26:171–87.

20. Akerman I, Tu Z, Beucher A, Rolando DMY, Sauty-Colace C, Benazra M, et al. Human pancreatic beta cell lncRNAs control cell-specific regulatory networks. Cell Metab. 2017;25:400–11.

21. Ding EL, Song Y, Manson JE, Hunter DJ, Lee CC, Rifai N, et al. Sex hormone-binding globulin and risk of type 2 diabetes in women and men. N Engl J Med. 2009;361:1152–63.

22. Peila R, Arthur RS, Rohan TE. Association of sex hormones with risk of cancers of the pancreas, kidney, and brain in the UK biobank cohort study. Cancer Epidemiol Biomarkers Prev. 2020;29:1832–6.

23. Peng H, Chen R, Brentnall TA, Eng JK, Picozzi VJ, Pan S. Predictive proteomic signatures for response of pancreatic cancer patients receiving chemotherapy. Clin Proteomics. 2019;16:31–42.

24. Ellingwood SS, Cheng A. Biochemical and clinical aspects of glycogen storage diseases. J Endocrinol. 2018;238:R131–R41.

25. Gentry MS, Roma-Mateo C, Sanz P. Laforin, a protein with many faces: glucan phosphatase, adapter protein, et alii. FEBS J. 2013;280:525–37.

26. Ward LD, Kellis M. HaploReg v4: systematic mining of putative causal variants, cell types, regulators and target genes for human complex traits and disease. Nucleic Acids Res. 2016;44:D877–81.

27. Hannukainen JC, Borra R, Linderborg K, Kallio H, Kiss J, Lepomaki V, et al. Liver and pancreatic fat content and metabolism in healthy monozygotic twins with discordant physical activity. J Hepatol. 2011;54:545–52.

28. Wojcik GL, Graff M, Nishimura KK, Tao R, Haessler J, Gignoux CR, et al. Genetic analyses of diverse populations improves discovery for complex traits. Nature. 2019;570:514–8.

29. Liu Y, Basty N, Whitcher B, Bell JD, Sorokin E, van Bruggen N, et al. Systematic quantification of health parameters from UK Biobank abdominal MRI using deep learning. BioRxiv. 2020.

30. Kolonel L, Henderson B, Hankin J, Nomura A, Wilkens L, Pike M, et al. A multiethnic cohort in Hawaii and Los Angeles: Baseline characteristics. American Journal of Epidemiology. 2000;151:346–57.

31. Lim U, Monroe KR, Buchthal S, Fan B, Cheng I, Kristal BS, et al. Propensity for intra-abdominal and hepatic adiposity varies among ethnic groups. Gastroenterology. 2019;156:966–75.

32. Sakai NS, Taylor SA, Chouhan MD. Obesity, metabolic disease and the pancreas-Quantitative imaging of pancreatic fat. Br J Radiol. 2018;91:20180267.

33. Chouhan MD, Firmin L, Read S, Amin Z, Taylor SA. Quantitative pancreatic MRI: a pathology-based review. Br J Radiol. 2019;92:20180941.

34. Park SL, Li Y, Sheng X, Hom V, Xia L, Zhao K, et al. Genome-Wide Association Study of Liver Fat: The Multiethnic Cohort Adiposity Phenotype Study. Hepatol Commun. 2020;4:1112–23.

35. Shepherd JA, Ng BK, Sommer MJ, Heymsfield SB. Body composition by DXA. Bone. 2017;104:101–5.

36. Le Marchand L, Wilkens LR, Castelfranco AM, Monroe KR, Kristal BS, Cheng I, et al. Circulating biomarker score for visceral fat and risks of incident colorectal and postmenopausal breast cancer: The Multiethnic Cohort Adiposity Phenotype Study. Cancer Epidemiol Biomarkers Prev. 2020;29:966–73.

37. Matthews D, Hosker J, Rudenski A, Naylor B, Treacher D, Turner R. Homeostasis model assessment: insulin resistance and fl-cell function from fasting plasma glucose and insulin concentrations in man. Diabetologia. 1985;28:412–9.

38. Mojiminiyi OA, Abdella NA. Effect of homeostasis model assessment computational method on the definition and associations of insulin resistance. Clin Chem Lab Med. 2010;48:1629–34.

39. Friedewald W, Levy R, Fredrickson D. Estimation of the concentration of low-density lipoprotein cholesterol in plasma, without use of the preparative ultracentrifuge. Clinical Chemistry. 1972;18.

40. Bien SA, Wojcik GL, Zubair N, Gignoux CR, Martin AR, Kocarnik JM, et al. Strategies for enriching variant coverage in candidate disease loci on a multiethnic genotyping array. PLoS One. 2016;11:e0167758.

41. The 1000 Genomes Project Consortium. An integrated map of genetic variation from 1,092 human genomes. Nature. 2012;491:56–65.

42. McCarthy S, Das S, Kretzschmar W, Delaneau O, Wood AR, Teumer A, et al. A reference panel of 64,976 haplotypes for genotype imputation. Nat Genet. 2016;48:1279–83.

43. Price AL, Patterson NJ, Plenge RM, Weinblatt ME, Shadick NA, Reich D. Principal components analysis corrects for stratification in genome-wide association studies. Nat Genet. 2006;38:904–9.

44. Polfus LM, Darst BF, Highland H, Sheng X, Ng MCY, Below JE, et al. Genetic Discovery and Risk Characterization in Type 2 Diabetes across Diverse Populations. (in press).

45. Willer CJ, Li Y, Abecasis GR. METAL: fast and efficient meta-analysis of genomewide association scans. Bioinformatics. 2010;26:2190–1.

